# Attenuation of muscle spindle firing with artificially increased series compliance during stretch of relaxed muscle

**DOI:** 10.1101/2023.05.08.539853

**Authors:** Emily M Abbott, Jacob D Stephens, Surabhi N Simha, Leo Wood, Paul Nardelli, Timothy C Cope, Gregory S Sawicki, Lena H Ting

## Abstract

Muscle spindles relay vital mechanosensory information for movement and posture, but muscle spindle feedback is coupled to skeletal motion by a compliant tendon. Little is known about the effects of tendon compliance on muscle spindle feedback during movement, and the complex firing of muscle spindles make these effects difficult to predict. Our goal was to investigate changes in muscle spindle firing using added series elastic elements (SEEs) to mimic a more compliant tendon, and to characterize the accompanying changes in firing with respect to muscle-tendon unit (MTU) and muscle fascicle displacements (recorded via sonomicrometry). Sinusoidal, ramp-hold-release, and triangular stretches were analyzed to examine potential changes in muscle spindle instantaneous firing rates (IFRs) in locomotor-and perturbation-like stretches as well as history dependence. Added SEEs effectively reduced overall MTU stiffness and generally reduced muscle spindle firing rates, but the effect differed across stretch types. During sinusoidal stretches, peak firing rates were reduced and IFR was strongly correlated with fascicle velocity. During ramp stretches, SEEs reduced the dynamic and static responses of the spindle during lengthening but had no effect on initial bursts at the onset of stretch. Notably, IFR was negatively related to fascicle displacement during the hold phase. During triangular stretches, SEEs reduced the mean IFR during the first and second stretches, affecting the history dependence of mean IFR. Overall, these results demonstrate that tendon compliance may attenuate muscle spindle feedback during movement, but these changes cannot be fully explained by reduced muscle fascicle length and velocity.

**New Findings:** *What is the central question of the study?:* Little is known about the effects of tendon compliance on muscle spindle function. We asked whether increasing the series compliance the muscle-tendon unit muscle spindle Ia responses to stretch. We also test the relationship between muscle spindle firing rates and muscle fascicle biomechanics.

*What is the main finding and its importance?:* Muscle spindle firing was generally attenuated with added series compliance, with the exception of the initial burst at the onset of stretch. Overall, the changes depended upon stretch profiles, and could not be fully explained by muscle fascicle length and velocity.

## Introduction

Muscle spindle sensory organs provide critical mechanosensory information for movement, but the coupling of sensory feedback to joint mechanics depends on mechanical properties of the muscle-tendon unit (MTU), which can change in aging and disease [17, 21, 33, 46]. Muscle spindle sensory feedback contributes to the control of movement and maintenance of posture [1, 19, 22, 25, 28, 40, 41]. Muscle spindle afferents fire reliably to stretch of the muscle-tendon unit, particularly when the muscle is in a relaxed state, and fire in relation to both the speed and extent of the movement. For example, during locomotion, the muscle is stretched cyclically at a relatively low frequency where muscle spindles exhibit approximately sinusoidal firing rates [24, 36-38]. In contrast, during discrete perturbations muscles can be very rapidly stretched, and muscle spindles exhibit very high and transient firing rates with various components that are sensitive to the acceleration, velocity, and/or displacement of the entire MTU [9, 13, 26, 42]. MTU stretch reflects the sum of the displacement of the muscle fascicle and the tendon that is mechanically in-series with the muscle. Thus, muscle spindle sensory organs throughout the belly of the muscle are only indirectly coupled to skeletal motion through the compliant tendon.

Discerning the effects of increased tendon compliance on muscle spindle firing during behavior in either humans or animals is extremely difficult. There is mounting evidence that tendons become more compliant in older adults [21, 24, 33, 46, 47]. Accordingly, older adults have decreased proprioceptive ability that affects their movement and posture [11, 14, 34, 45]. However, aging studies cannot isolate the effects of tendon compliance from other physiological effects of age, such as changes to the muscular and nervous systems, and directly manipulating tendon compliance *in vivo* is highly invasive and could alter sensorimotor control strategies. Most recordings from animal muscle spindle afferents occur in acute, anesthetized preparation such as the rat, where MTU stretch is directly controlled and muscle spindle afferent firing is recorded from the dorsal roots of the spinal cord. Prior work has examined the role of tendon compliance on muscle spindle firing but examined steady firing rates at different lengths rather than over the course of MTU stretch, and did not directly record fascicle lengths [7]. Acute studies in aged rats do show that the muscle tendon unit must be stretched farther to initiate firing of action potentials [31], however age affects both tendon and extracellular matrix compliance, and changes in animal models are different than in humans [18], limiting direct comparisons. Therefore, our goal was to directly manipulate series compliance and to observe the immediate effects on muscle spindle firing during stretch.

Prior interpretations of muscle spindle firing behavior have largely relied on its association with measurements of the MTU, rather than measurements of the muscle fascicle. However, forces and displacements at the level of the spindle are indirectly coupled to the motion of the skeleton via the series elasticity of the tendon [12, 23]. Further, the forces carried by the tendon are shared between the muscle fascicles and the extracellular matrix such that the fascicle displacement cannot be easily inferred from MTU mechanics. Evidence from measures of human muscle spindle firing shows a close relationship to muscle fascicle stretch in slow, postural-sway like stretches [6]. However, the relationships between spindle afferent firing responses with respect to muscle fascicle biomechanics is still an open question.

Given the complexities of muscle spindle firing, the effect of tendon compliance on muscle spindle firing is also likely to be complex and may depend on the specific stretch conditions tested. Muscle spindle firing during locomotion in cats is approximately sinusoidal, resembling MTU velocity [36, 37]. However, muscle spindle firing during passive sinusoidal stretches of the MTU at low frequencies in the anesthetized cat have been shown to follow MTU length in time, within nonlinear scaling to stretch amplitude [16, 27]. In more rapid stretches, muscle spindles exhibit even more complex firing patterns. Initial bursts consist of very rapid firing at stretch onset and are attributed to either stretch acceleration or rate change in force, i.e. yank [2-4, 15, 39, 42, 43]. During ramp stretches the firing rate is substantially increased with respect to muscle length change [10]. When held at a new, longer, length, muscle spindle firing rates decrease rapidly and then slowly relax to a new steady-state firing rate [5]. Finally, these features of firing do not have a unique relationship to stretch features, as repeated triangle stretches reveal an absence of the initial burst and lower dynamics response in the second stretch; this history dependence has been attributed to nonlinearities in muscle stiffness [2-4, 15, 32, 39].

Here we hypothesized that increased tendon compliance (or decreased stiffness) would reduce muscle spindle firing during stretch. We further hypothesized that muscle spindle firing would be more closely related to muscle fascicle versus MTU stretch. We experimentally increased tendon compliance and to characterize the effects on muscle spindle afferent firing during MTU stretch of relaxed muscle while recording muscle spindle firing and muscle fascicle length. Effective tendon compliance in anesthetized animals was reduced by adding a series elastic elements (SEEs) to the end of the MTU. The same sinusoidal, ramp-hold-release, and repeated triangular stretches were applied to the MTU and to the MTU+SEE complex. Simultaneously, we recorded muscle spindle firing from identified Ia afferents, muscle fascicle length using sonomicrometry, and MTU force. Given the nonlinear relationship between MTU stretch amplitude and muscle spindle firing in sinusoidal stretches, we tested whether potential changes in muscle spindle firing could be accounted for by changes in fascicle displacement and velocity between control and SEE groups. In ramp-hold-release stretches, we compared the relationships of muscle spindle firing during the hold phase to fascicle displacement and force, as the relaxation of muscle spindle IFR during the hold phase has been speculated as being driven by fascicle length changes. In triangular stretches, we tested whether the stretch response after the first stretch could be attributed to changes in fascicle displacement or MTU force.

## Methods

### Animal Care

All procedures and experiments were approved by the Georgia Institute of Technology Institutional Animal Care and Use Committee. Adult female Wistar rats (250–300 g; Charles Rivers Laboratory, Wilmington, MA) were studied in terminal experiments only and were not subject to any other experimental procedures. All animals were housed in clean cages and provided food and water ad libitum in a temperature and light controlled environment in the Georgia Institute of Technology Physiological Research Laboratory (protocol number A18042).

Surgery was carried out in the same manner described in previous experiments from this laboratory [48]. In brief, rats were deeply anesthetized (complete absence of withdrawal reflex) by inhalation of isoflurane, initially in an induction chamber (5% in 100% O_2_) and for the remainder of the experiment via a tracheal cannula (1.5–2.5% in 100% O_2_). Vital signs were continuously monitored including, core temperature (36-38 °C), PCO2 (3-5%), respiratory rate (40–60 breaths/min), pulse rate (300-450 bpm) and SPO2 (>90%). Anesthesia level and vital signs were maintained by adjusting isoflurane concentration, radiant and water-pad heat sources, and by scheduled subcutaneous injection of saline (1ml/hour subcutaneous). Surgical preparation followed by data collection lasted up to 8 hours.

### Data collection

To examine the effects of tendon compliance on muscle spindle behavior, we added elastic elements to the end of the medial gastrocnemius MTU to increase the series compliance of the MTU, mimicking a more compliant tendon. In SEE conditions, “MTU” refers to the MTU and SEE complex as one unit. Sinusoidal stretches were applied at 2 mm amplitude at 2 Hz, ramp-hold-release stretches at 3 mm length change at a velocity of 20 mm/s, and triangular stretches consisted of 3 repetitions of 3 mm length changes at 3.5 mm/s. MTU length and force were measured from a servomotor (AURORA MOTOR 310C-LR), and muscle fascicle length was measured with sonomicrometry crystals (Sonometrics) implanted along the same muscle fascicle within the MG, all sampled at 17.8 kHz (Fig. 1B). Recordings from spindle afferents were taken intracellularly via glass microelectrode inserted into dorsal root afferents. Ventral root motor efferents were also stimulated during these experiments, however the current analysis only considers passive stretches. Sinusoidal stretches incorporated ventral root stimulation following several passive cycles, and only the initial passive cycles are analyzed here. Data was collected and stored using Cambridge Electronic Design (CED) Power 1401 and Spike2 software and exported to MATLAB for analysis. Statistical comparisons were performed using R software.

**Figure 1.**
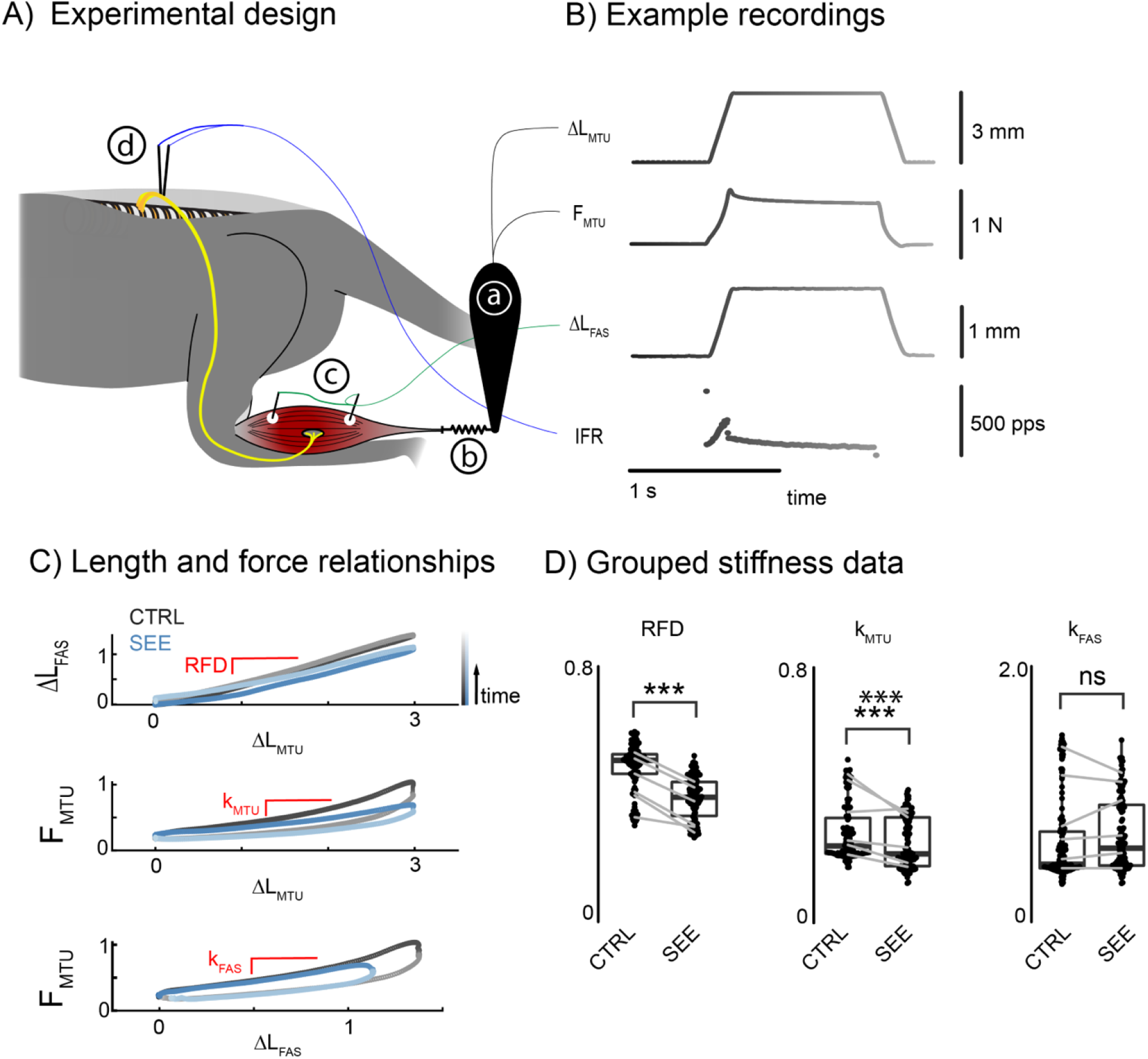
Experimental design, sample data, and stiffness analysis with added SEE. A) The medial gastrocnemius (MG) of anesthetized rats was dissected away from the rest of the triceps surae and attached to a length-controlled servomotor (a). Added SEEs were tethered between the end of the Achilles tendon at the calcaneous and the arm of the motor to effectively increase the series compliance of the MTU (b). Muscle spindle afferent action potentials were recorded intracellularly from the dorsal roots of the spinal cord (c), and sonomicrometry probes were implanted along a muscle fascicle of the MG to record the length change of muscle fascicles during stretch (d). B) Example recordings of MTU length change, MTU force, fascicle length change, and muscle spindle IFR during ramp-hold-release stretches. C) Examples of fascicle displacement versus MTU displacement, MTU force versus MTU displacement, and MTU force vs fascicle displacement. Line colors correspond to time in order to display differences in lengthening and shortening, corresponding to the shading in (B). Labelled red lines represent the metrics analyzed in (D). D) Grouped metrics show reduced relative fascicle displacement (RFD), reduced MTU stiffness (k_MTU_), and fascicle stiffness (k_FAS_) at 0.5 N in ramp stretches. Box plots indicate the mean and first and third quartiles, gray lines indicating change in mean for each animal. *** indicate p<0.001.

### Data Processing

Sonomicrometry readings were calibrated by recording channel voltage and measuring the distance between the probes in the muscle at resting length and at 1mm of MTU displacement with no series elastic element. Motor position and force were also recorded at resting length, and these values were used as baseline values for MTU length and force. Measurements of motor position, motor force, and fascicle displacement were first down-sampled by a factor of 20 to approximately 900 Hz, then lowpass filtered with a 4th-order Butterworth filter with an 80 Hz cutoff. Then data were smoothed and differentiated with a 2nd-order Savitsky-Golay filter with a window width of 41 samples (approximately 46 ms). Sonomicrometry measurements were calibrated by measuring the length between the two crystals along the muscle fascicle corresponding to two measurements, in order to match voltage values from the data channel to physical lengths. The calibration values were collected from all experiments, and the mean scaling factor was used as the scaling factor for all trials.

### Data Analysis

To validate the effects of SEEs, the length change of the fascicle relative to the MTU, the stiffness of the MTU, and the stiffness of the fascicle were computed for each stretch type. Stiffness was estimated by fitting a 2^nd^-order polynomial to force/length curves at 0.5 N in ramps and triangles, and 0.4 N in sinusoids, and computing the slope of the tangent line.The relative fascicle displacement (RFD) was estimated in the same manner, estimating the slope of the tangent line (dL_Fas_/dL_MTU_) at 0.5 or 0.4 N.

During sinusoidal stretches, peak firing rate and mean firing rate were analyzed following the first cycle to analyze changes in spindle responses at a steady state and ignore history dependent spindle firing. During ramp-hold-release stretches, initial bursts, dynamic responses, and static responses were extracted from firing rates (Fig. 3C). The static response was taken as the firing rate approximately 0.5 s into the hold phase. During triangular stretches, mean firing rate and spike counts were computed for the first and second stretches, along with initial burst. History dependence was analyzed by comparing the differences in mean IFR and spike count between the first and second stretches for each trial. Linear regression analyses were performed only after the first stretches in sinusoids and triangles, and only on the static response in ramp-hold-release stretches, as forces and length measurements are not expected to predict the more dynamic components of firing responses.

Trials were excluded according to IFR and force recordings. Trials with a lack of reliable spiking, considerable aberrant firing, or lack of initial bursts in ramp-hold-release or triangular stretches were excluded. Additionally, trials with considerable noise in force recordings were excluded to avoid inaccurate stiffness estimates.

Statistical comparisons of firing and stiffness were conducted with linear mixed models in R, using the *lmer* functionality of the *lmerTest* package [20]. In stiffness and fascicle displacement analyses, the effects of tendon compliance were determined by linear mixed models, with individual animals serving as random effects and SEE as a fixed effect. In firing rate comparisons, individual afferents served as random effects and SEE as a fixed effect. Linear regressions were computed in MATLAB, and per-trial R^2^ values were exported and analyzed in R. Reported values are the mean ± SD unless stated otherwise.

## Results

Overall, adding the series elastic element reduced fascicle displacement relative to MTU displacement during stretch and reduced effective tendon stiffness (Fig. 1C). Relative fascicle displacement was reduced in the SEE condition by 0.085 ± 0.004 mm/mm (mean ± SE) in ramps (p <1 e-14, 306 trials across 6 animals), 0.06±0.02 mm/mm in triangles (p<1e-4, 78 trials across 5 animals), and 0.067±0.01 mm/mm in sinusoids (p<1e-7, 85 trials across 5 animals). MTU stiffness was reduced by 0.040±0.004 N/mm (p<2e-16) in ramps, 0.031±0.015 N/mm in triangles (p=0.043), and 0.032±0.0050 N/mm (p<2e-7) in sinusoids. In contrast, there were no differences in fascicle stiffness with added SEE across condition (all p>0.13)

During sinusoidal stretches, adding the SEE reduced muscle spindle firing rates (see exemplar in Fig. 2A). Although there was no reduction in mean firing rate with added SEE (p>0.05, 85 trials across 10 afferents), peak firing rate was reduced by 9.8±4.2 pulses per second (pps) (p=0.022) (Fig. 2C). In some cases, greater firing responses were observed in the first cycle (Fig. 2). IFR was generally poorly correlated with muscle fascicle length after the first stretch (R^2^ = 0.05 ± 0.05), and the regression slope was reduced in SEE conditions by 22.2% (5.40 ± 2.16 pps/mm, mean ± SE, p = 0.0144). IFR was also weakly correlated with fascicle velocity (R^2^ = 0.11 ± 0.09), but the regression slope was increased by 65.1% in SEE conditions (control rate = 2.47 ± 0.37, change = 1.61±0.21 pps/mm/s, mean ± SE, p<1e-10). Additionally, MTU force was weakly correlated with muscle spindle firing rate (R^2^ = 0.12 ± 0.11) but did not show a difference in regression slope (p = 0.814) or intercept (p = 0.761) between control and SEE conditions.

**Figure 2.**
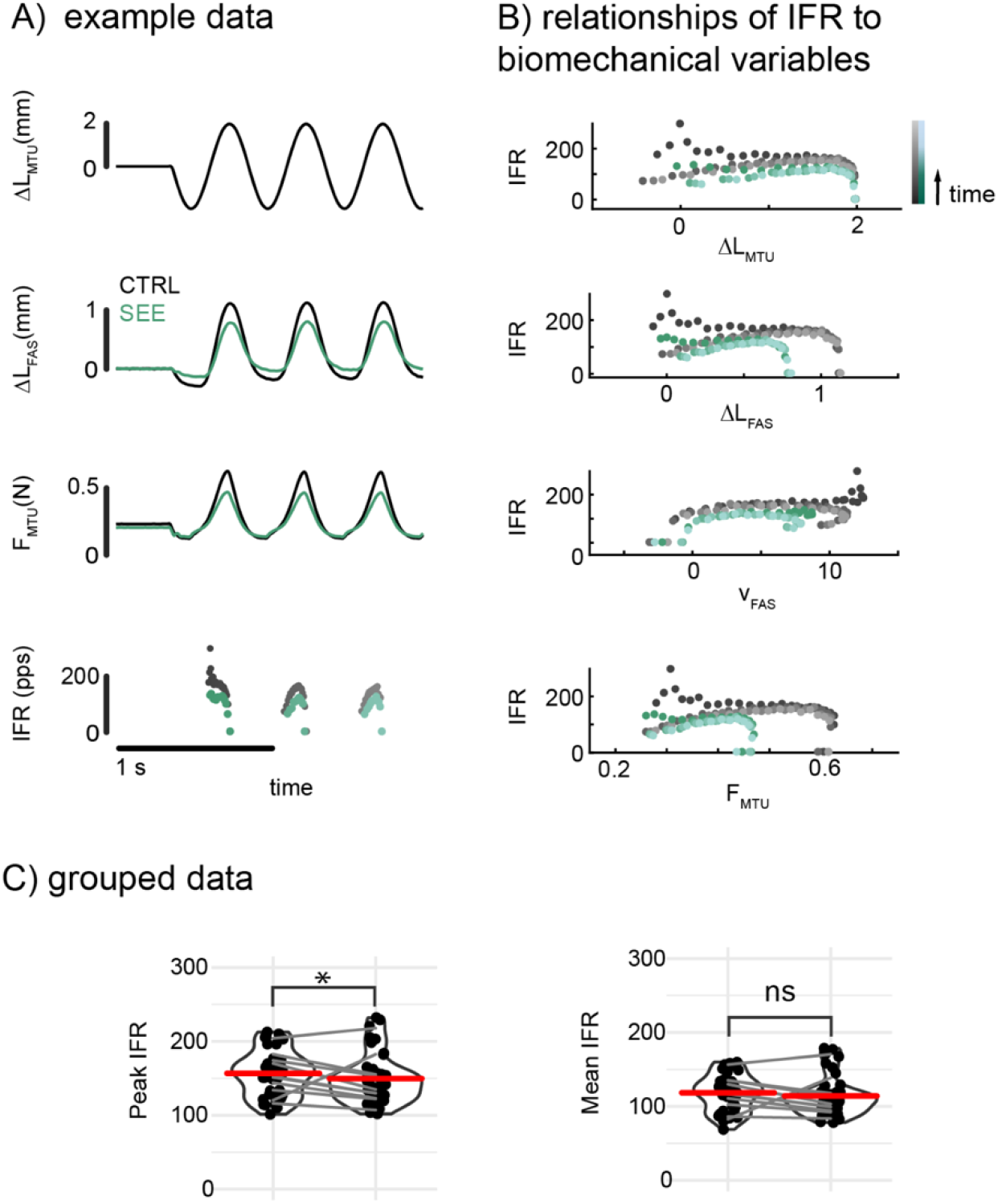
Changes in firing rates and relationships to muscle and MTU mechanics during 2 mm, 2 Hz sinusoidal stretches of the MTU. A) Addition of SEE revealed reduced fascicle displacement in both shortening and lengthening, reduced MTU force, and reduced muscle spindle IFR (control-black, SEE-green, darker points appearing earlier in time). B) Relationships between IFR and MTU displacement, fascicle displacement and velocity, and MTU force reveal the strongest relationship between IFR and MTU force (R^2^= 0.12 ±0.11). Initial burst is evident in both control and SEE. C) Statistical comparisons of peak and mean IFR after the first cycle reveal a significant reduction in peak IFR of 9.8 ± 4.2 pps (p = 0.022), but no reduction in mean IFR (p > 0.05) with added SEE. * indicate p<0.05, ns – no significance

During ramp-hold-release stretches, adding the series elastic element did not alter the initial burst at the onset of firing, but firing during the ramp and hold were decreased. Fascicle length changes were similar to MTU changes, but with reduced amplitude with added SEE (see Fig. 3A for example). The MTU force rose faster than the ramp displacement and gradually relaxed during the hold period. At the onset of the ramp, muscle spindle firing exhibited an initial burst that was not changed by the added SEE (control rate= 329±115, SEE rate = 321±92, p>0.13). The dynamic response of the muscle spindle during the ramp phase (3 mm, 20 mm/s), was attenuated with SEE by 28.0±1.8 pps (p < 2e-16). The static firing rate during the hold phase also decreased with added SEE by 2.3±1.1 pps (p<0.03). Accordingly, the dynamic index, indicating the decrease in firing rate during the hold phase, was also reduced with added SEE by 25.7±1.4 pps (p < 2e-16).

**Figure 3.**
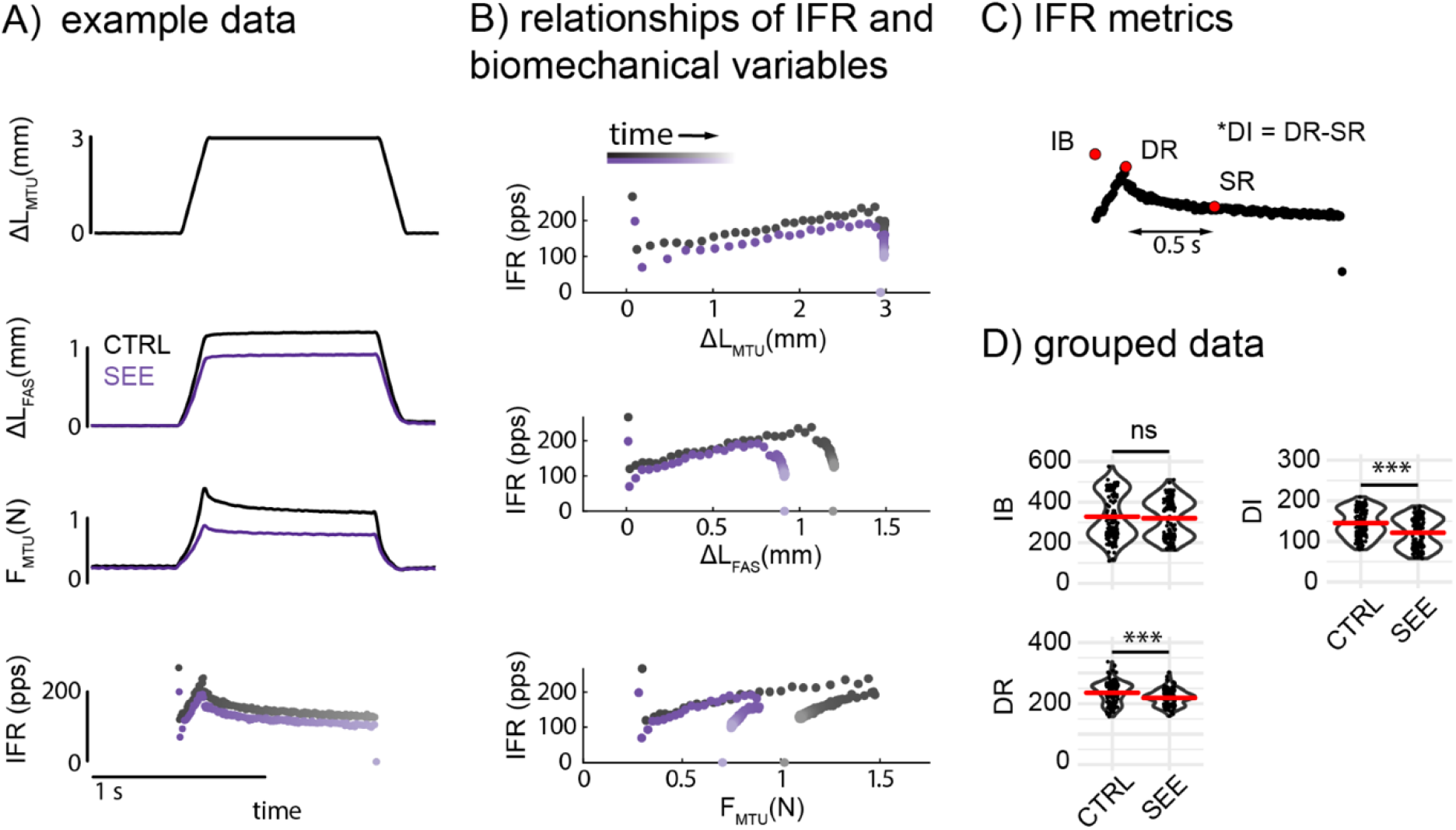
IFR vs muscle and MTU mechanics during ramp-hold-release stretches. 3 mm, 20 mm/s stretches were applied to the MTU, followed by a > 1s hold phase and subsequent release. A) Muscle fascicle displacement and MTU force were subsequently reduced with added SEE (black-CTRL, purple-SEE). B) Relationships between IFR and MTU displacement, fascicle displacement, and MTU force reveal none of these variables predict initial burst or dynamic response, but MTU force shows a concomitant decrease with the decrease in IFR during the hold phase. Lines indicate zero-intercept regression slope and shading corresponds to time as in panel A. C) Examples of initial burst (IB), dynamic response (DR), static response (SR) measured 0.5 s after the DR, and dynamic index (DI) as the difference in the dynamic and static responses. D) Across trials, SEE resulted in a 28.0 ± 1.8 pps decrease in DR (p < 2e-16) and 25.7 ± 1.4 pps decrease in DI (p < 2e-16) with no change in IB (p>0.13). *** indicate p<0.001, ns – no significance

During the hold phase, muscle spindle IFR was strongly correlated with MTU force (R^2^ = 0.67 ± 0.19), and weakly correlated with fascicle displacement and velocity and MTU displacement and velocity (all mean R^2^ between 0.22 and 0.35). During the hold phase, muscle fascicle displacement did not predict the observed changes in muscle spindle firing rate. As muscle spindle firing decreased, the muscle fascicle lengthened (Fig. 3B) and linear regression analysis revealed a negative relationship between IFR and fascicle displacement in both control (slope = -1317 ± 1349, R^2^ = 0.29 ± 0.23, 156 trials) and SEE trials (slope = -1217 ± 943, R^2^ = 0.40 ± 0.21, 150 trials). Positive relationships were found between IFR and MTU force in control (slope = 422 ± 224, R^2^ = 0.71 ± 0.19) and SEE trials (slope = 581 ± 392, R^2^ = 0.64 ± 0.19), however the slopes of these fits were different (p<2e-16).

During repeated triangular stretches, SEE increased the history dependence of muscle spindle mean IFR. At the onset of the first stretch, muscle spindles produced initial bursts in both conditions as well as a response during the ramp stretch. In the second and subsequent stretches, the initial burst was absent, and the response during the ramp stretch was reduced. During the first stretch, SEE had no effect on initial bursts or spike counts but did reduce the mean IFR by 6.5±2.6 pps (p = 0.016). In the second stretch, SEE similarly did not affect spike counts but did reduce the mean IFR by 9.1±2.1 pps (p < 4e-5). In the same manner, the history dependence of spike counts was not affected by SEE, but the history dependence of mean IFR was increased with added SEE by 2.8±2.1 pps (p < .01).

During the second and third stretches, muscle spindle IFR was moderately correlated with MTU force (R^2^ = 0.52 ± 0.09), MTU displacement (R^2^ = 0.49 ± 0.13), and fascicle displacement (R^2^ = 0.47 ± 0.15), and not correlated with MTU nor fascicle velocity (R^2^ = 0.03±0.05 and 0.08±0.09, respectively). However, overall, the relationships between muscle spindle IFR with respect to all of the biomechanical variables were not unique and changed between the first and second stretches, and with added SEE (Fig 4B).

**Figure 4.**
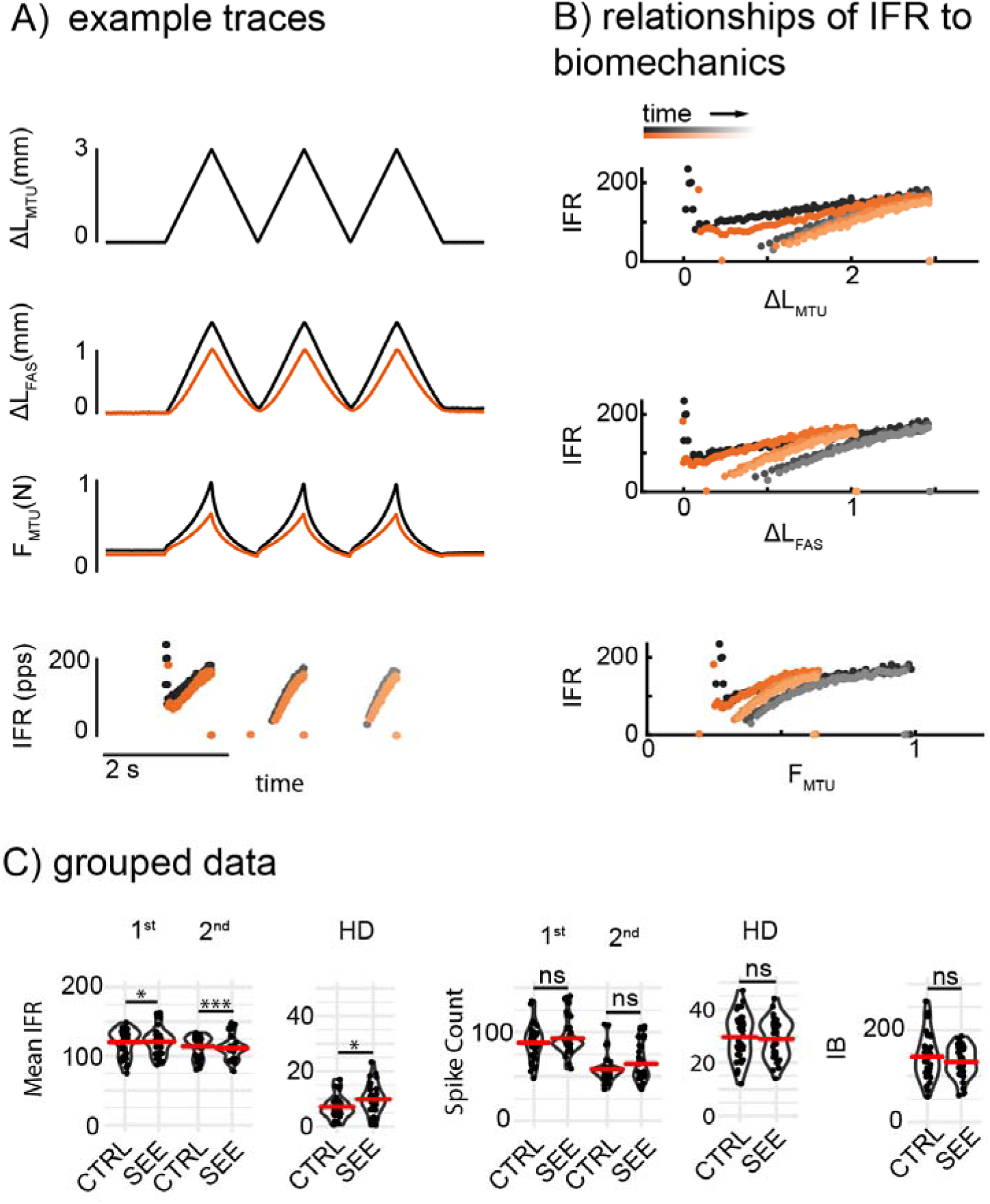
Relationships of MTU displacement and force and fascicle displacement to muscle spindle IFR during triangular stretches. Effects of series elasticity on muscle fascicle displacement, MTU force, and muscle spindle IFR repeated triangle stretch trials of 3 mm, 3.5 mm/s. A) Muscle fascicle displacement and MTU force were reduced with added SEE (black-CTRL, orange-SEE). In both conditions, stretches evoked IBs and DRs in the first stretch (black-CTRL, dark orange-SEE) that were absent in subsequent stretches (gray-CTRL, light orange-SEE). B) Relationships between IFR and MTU displacement, fascicle displacement, and MTU force. All variables reasonably predict IFR during the second and third stretches, but not the IB or DR of the first stretch. Lines indicate zero-intercept regression slope. C) Across groups, SEE resulted no change in IB (p > .05) or spike counts during the first or second stretch, and subsequently spike count history dependence. However, SEE did result in a 6.5 ± 2.6 pps decrease in mean IFR during the first stretch (p = 0.016) and a 9.1 ± 2.1 pps decrease in mean IFR in the second stretch (p < 4e-5), thus increasing mean IFR history dependence by 2.8 ± 2.1 pps (p < .01). * indicate p<0.05, *** indicate p<0.001, ns – no significance

## Discussion

Our data show that increasing the effective compliance of tendon decreased the firing rate in muscle spindles during muscle stretch. Reductions in firing rates with added seried elasticity depended on the muscle stretch profile, with reduced peak firing rates in sinusoidal stretches and in the ramp and hold phases of discrete stretches. These data suggest that lower tendon stiffness could reduce sensory feedback in locomotion and movement. However, the lack of change in initial bursts at the onset of stretch indicate that the considerable sensitivity of muscle spindles to small changes in length at the onset of stretch may not be significantly impacted by tendon compliance. Thus, lower tendon stiffness may not adversely affect sensory feedback for perturbation detection. Notably, although muscle spindle firing was related to fascicle displacement and velocity during continuous stretches (sinusoids, second and later triangles), it had non-unique relationships to fascicle displacement during ramp stretches, and negative relationships to fascicle length during the hold phase after a ramp. Our data show that fascicle length does not completely explain the nuances of muscle spindle behavior, and that more complex models may be necessary to understand how changes in mechanical properties of connective tissues affect sensory feedback from muscle spindles.

The experiments presented here validated the use of adding series springs to experimentally increase tendon compliance and decouple biomechanical relationships within the muscle-tendon unit. Using sonomicrometry revealed that the relative stretch in muscle fascicles with respect to the effective MTU was reduced, indicating increased stretch in the effective tendon. Further, total muscle fascicle stiffness remained unchanged while total MTU stiffness was reduced when adding SEEs. At longer stretch lengths, the difference in the MTU force and length was larger, as the nonlinear elasticity of the muscle tissues was engaged. Thus, adding tendon compliance changes the distribution of length (and force via stiffness) within the compartment of the MTU enabling experiments to probe the role of different biomechanical signals (e.g. length, velocity; force, yank) in shaping muscle spindle output. A limitation of the current experiment is that, in the added SEE condition, *both* MTU force and fascicle length decreased, making it difficult to clearly distinguish the roles of force vs length-related dynamics as driving inputs of muscle spindle firing. In sinusoidal stretches, muscle spindle firing was not well-correlated with any of the tested biomechanical variables, and some relationships differed between control and SEE trials. Contrary to prior work [6], muscle fascicle displacement and velocity were not well correlated with muscle spindle firing during sinusoidal stretches. However, this prior work incorporated phase shifts between the measured IFR and displacement and velocity which were not included here. While correlations were not strong, the change in regression coefficients is still indicative of a more complex relationship to muscle fascicle biomechanics than simply displacement or velocity. Further, while MTU force showed a poor correlation, there was no evidence of an effect of added compliance (SEE) on this relationship. However, the relationships of IFR to any of these variables is markedly complex (Fig. 2B), thus a more rigorous model comparison is warranted to further test the relationships of muscle spindle firing to fascicle mechanics.

As the initial burst of muscle spindles has been hypothesized to play a role in postural control [20, 49, 50], it seems that tendon compliance may not significantly impact postural control responses, as the high sensitivity of muscle spindles to small stretches is retained. The lack of change in initial bursts in either ramp-hold-release or triangular stretches may be due to the added compliance having less of an effect at short MTU lengths. Initial bursts of muscle spindles have previously been related to the rapid increase in force on the muscle spindle at the onset of stretch due to intrafusal muscle spindle fiber short-range stiffness [2, 3, 10, 32, 39]. As the effect of a series elastic element on MTU force is most pronounced at longer MTU displacements (Figs. 1-4), it may not reduce the rate at which force is applied to the spindle particularly at the onset of stretch, thus preserving initial bursts with increased tendon compliance.

The negative relationships between muscle spindle firing and fascicle length at constant MTU length may be due to the stress relaxation of extrafusal skeletal muscle fibers and intrafusal muscle spindle fibers after being stretched. It has been suggested that the slowing of muscle spindle firing during the hold phase is due to fascicle shortening, however fascicle displacement *increased* during the hold phase of ramp-hold-release stretches as firing decreased (Fig. 3B). Prior model predictions also showed that changes in muscle fascicle length are in opposition to muscle spindle firing rates during conditions where MTU length does not predict muscle spindle firing [3]. While the relationship between IFR and MTU force was different in SEE trials, it did predict the concomitant decrease in IFR. It has been suggested that the relaxation of muscle spindle firing during the hold phase is due to the stress relaxation of intrafusal muscle spindle fibers [5, 32], and both extrafusal and intrafusal fibers relaxing simultaneously. Thus, as intrafusal fibers relax, muscle spindle firing decreases, and as extrafusal fibers relax, MTU force decreases and fascicle length increases.

More complex modeling may be necessary to better understand the relationships between multiscale mechanics in the muscle and muscle spindle firing rate. While muscle spindle firing appears more force-related than fascicle-displacement-related during the hold phase, MTU force was only moderately correlated with MTU force here, and the relationship changed between control and SEE trials. Prior models relating MTU force and yank to muscle spindle firing rates in rats and cats [2, 3] are not suited to this data. In rats, the force contribution of extramysial tissue in the muscle has to be estimated from the displacement of the MTU and subtracted from the total MTU force. However, as MTU displacement measurements in these experiments also encompassed the added elastic measurement, it cannot be used as an input to an extramysial tissue model. It is not known if fascicle measurements from sonomicrometry can accurately predict extramysial force, as it may not measure the displacement of the connective tissues carrying the loads [8, 29, 30]. Further, this data fitting model is only accurate to the degree that extrafusal and intrafusal forces are approximately proportional. During triangle stretches, the similarity in muscle spindle firing in the second and third stretch is likely due to the intrafusal muscle going slack within the extrafusal muscle [2-4, 35]. Thus, it may be necessary to have a biophysical model to separately predict intrafusal and extrafusal muscle dynamics [2] to more accurately predict muscle spindle firing.

Changes in tendon compliance with aging and disease may have varied and context-dependent effects on muscle spindle sensory feedback. Increased tendon compliance generally reduced muscle spindle firing, but in complex relationships to muscle mechanics. The changes in muscle spindle firing likely depend on the nature of the movement, in addition to nonlinear properties of the muscle and tendon tissues. Further, the different relationships between muscle fascicle and muscle biomechanics across all stretch types suggest that muscle spindles do not simply encode muscle fascicle length.

## Author Contributions

Dr. Emily M Abbott – conception and design, data acquisition and analysis, drafting and revising, Jacob D. Stephens – data analysis and interpretation, drafting and revising, Dr. Surabhi N. Simha – data interpretation, drafting and revising, Leo Wood – data analysis, drafting and revising, Paul Nardelli: data acquisition, drafting and revising, Dr. Timothy C. Cope – conception and design, drafting and revising, Dr. Gregory S. Sawicki – conception and design, data analysis and interpretation, drafting and revising, Dr. Lena H. Ting – data analysis and interpretation, drafting and revising

All authors have approved the final version of this manuscript and agree to be accountable.

## Data availability statement

Data are available on request

